# The dynamic interaction of systemic inflammation and the hypothalamic-pituitary-adrenal (HPA) axis during and after major surgery

**DOI:** 10.1101/2021.08.16.456512

**Authors:** Daniel Galvis, Eder Zavala, Jamie J. Walker, Thomas Upton, Stafford L. Lightman, Gianni D. Angelini, Jon Evans, Chris A. Rogers, Kirsty Phillips, Ben Gibbison

**Author notes:** These authors contributed equally to this work (Daniel Galvis, Eder Zavala). (E.Z), (B.G.).

## Abstract

Major surgery and critical illness produce a potentially life threatening systemic inflammatory response. The hypothalamic-pituitary-adrenal (HPA) axis is one of the key physiological systems that counterbalances this systemic inflammation through changes in adrenocorticotrophic hormone (ACTH) and cortisol. These hormones normally exhibit highly correlated ultradian pulsatility with an amplitude modulated by circadian processes. However, these dynamics are disrupted by major surgery and critical illness. In this work, we characterise the inflammatory, ACTH and cortisol responses of patients undergoing cardiac surgery and show that the HPA axis response can be classified into one of three phenotypes: single-pulse, two-pulses and multiple-pulses dynamics. We develop a mathematical model of cortisol secretion and metabolism that predicts the physiological mechanisms responsible for these different phenotypes. We show that the effects of inflammatory mediators are important only in the single-pulse pattern in which normal pulsatility is lost – suggesting that this phenotype could be indicative of the greatest inflammatory response. Investigating whether and how these phenotypes are correlated with clinical outcomes will be critical to patient prognosis and designing interventions to improve recovery.

## INTRODUCTION

Major surgery and critical illness elicit a systemic inflammatory response of both clinical and scientific relevance^1^, which when uncontrolled leads to major morbidity and/or death^2,3^. One of the key physiological systems that regulates the inflammatory response in humans is the hypothalamic-pituitary-adrenal (HPA) axis, which controls the rhythmic secretion of adrenocorticotrophic hormone (ACTH) and cortisol. The dynamics of these hormones are transiently disrupted by stressors, with HPA axis activity following major surgery^4^ and critical illness^5^ constituting an important biological marker of the patient’s response. Many of the models for the HPA axis and inflammation used in clinical and scientific practice are based on linear relationships. Both inflammation and the HPA axis are cascades, with individual components having multiple sites of action in other systems and feedback on their own^6^. The secretion and effects of these hormones are also non-linear, making the overall effect at the level of an organism difficult to elucidate. Understanding how ACTH and cortisol are controlled during acute systemic inflammation (e.g. major surgery and critical illness) is vitally important because both excess and deficiency of these hormones are associated with death and major morbidity for patients^7^.

Under normal physiological conditions, ACTH and cortisol are in a state of continuous dynamic equilibration^8^ and exhibit highly-correlated ultradian pulsatility with an amplitude modulated by upstream circadian processes (Fig. 1A). ACTH and cortisol dynamics are disrupted after major surgery: mean concentrations of cortisol are higher, mean concentrations of ACTH are lower, and the correlation between ACTH and cortisol is altered^4^. A combination of animal physiology experiments and mathematical modelling has also shown that acute inflammation can disrupt the high correlation between ACTH and cortisol, a phenomenon called *dynamic dissociation^9^*. However, the interaction between the HPA axis and the inflammatory cascade in humans is unknown (Fig. 1B), in part because the lack of high-resolution time-course data has precluded accurate modelling.

**Figure 1.**
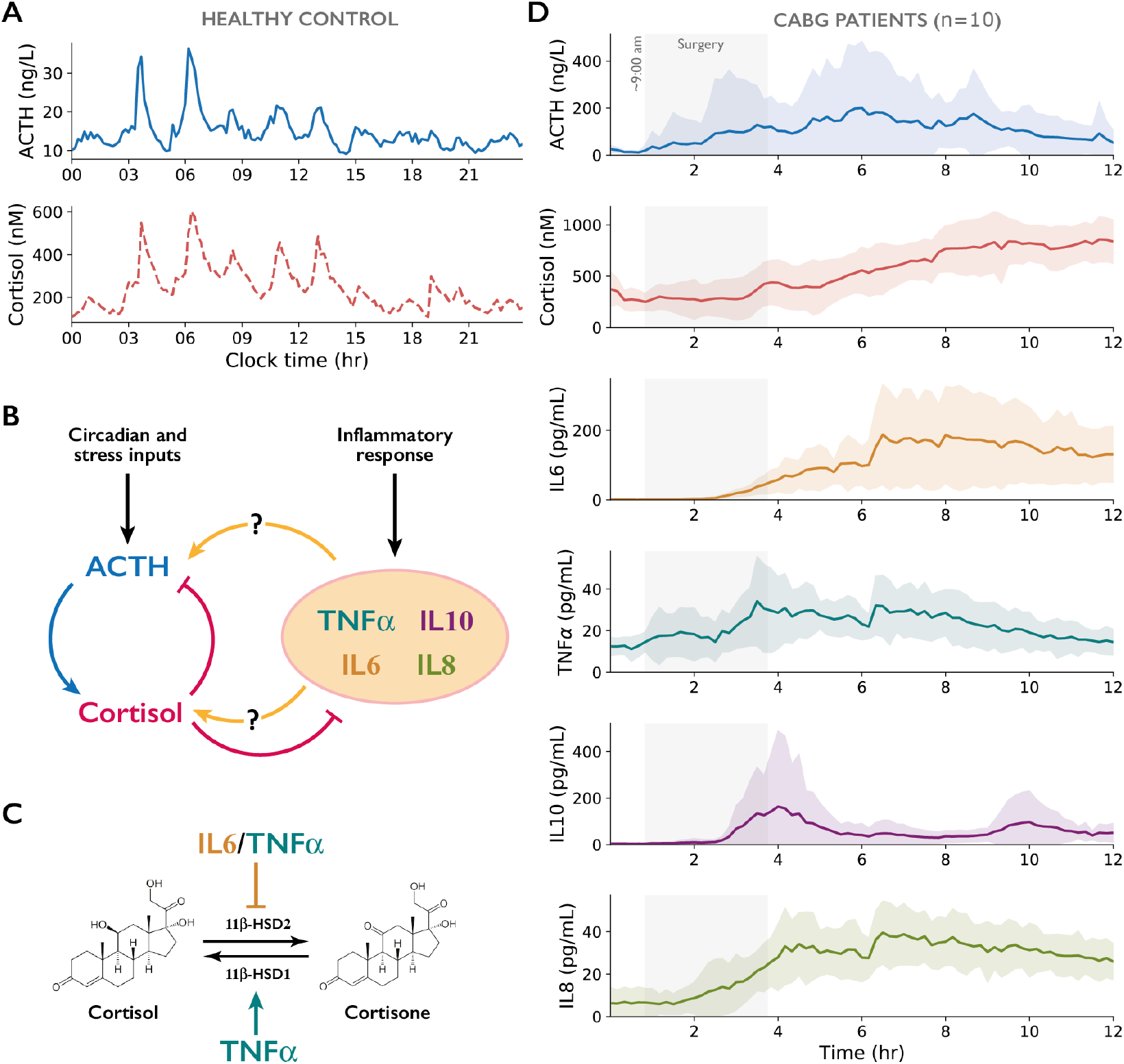
Dynamic stress and inflammatory responses following CABG. **A)** 24h ACTH and cortisol dynamic profile in a healthy control^4^. **B)** Systems-level interactions between the HPA axis and inflammatory mediators. **C)** Tissue-level regulation of cortisol availability by inflammatory mediators targeting 11β-HSD1/2. **D)** 12h dynamic profile (μ ± σ, n=10) of ACTH, cortisol, and inflammatory mediators during and after CABG. Grey shaded areas indicate the mean time span of CABG surgery. Detailed individual profiles are shown in Fig. S2.

The most likely mediators of the interactions between inflammation and the HPA axis are cytokines such as IL1α, IL6 and TNFα, and the glucocorticoid hormone cortisol. The isolated effects of cytokines on the HPA axis at the cellular level are well known (e.g. IL6 increases cortisol synthesis^10,11^). However, this information is not informative at the systems level due to the bi-directional feedback that all components exert on each other, in addition to the dynamic regulation of cortisol in target tissues (Fig. 1C). To address the question of how the inflammatory and HPA axis response interact at a systems level, we performed high-frequency serial blood sampling during and after coronary artery bypass grafting (CABG) surgery to generate profiles of ACTH, cortisol and inflammatory mediators IL6, IL8, IL10 and TNFα (Fig. 1D). These profiles were statistically classified according to dynamic phenotype and used to calibrate a mathematical model to characterise the underlying changes in HPA physiology.

## RESULTS

### Patient Characteristics

Ten patients underwent isolated CABG surgery via median sternotomy (See Table 1). CABG can take place with or without cardiopulmonary bypass (CPB). Five cases had their surgery performed with the use of CPB and five did not. When CPB was used, the duration was 67 ± 20.7 minutes. Patients and controls were not matched for age, height and weight, but were broadly similar in these parameters. No patient suffered major complications and this was indicated in the median critical care and hospital length of stay, which were shorter than the UK median (3.2 and 7.2 days respectively)^12,13^.

**Table 1.** Demographic and operative data of study participants.

|  | Controls (n=3) | Patients (n=10) |
| --- | --- | --- |
| Age | $54 \pm 13.8$ years | $65 \pm 6.2$ years |
| Height | $180 \pm 6.7$ cm | $173 \pm 6.4$ cm |
| Weight | $87.5 \pm 13.7$ kg | $83 \pm 10.7$ kg |
| BMI | $26.9 \pm 2.1$ kg/m<sup>2</sup> | $27.8 \pm 4.1$ kg/m<sup>2</sup> |
| Sampling duration | 24 hrs | 12 hrs |
| Anaesthesia start time | – | 8:26 AM ($\pm 22$ min) |
| Surgery start time | – | 9:19 AM ($\pm 22$ min) |
| Surgery end time | – | 12:15 PM ($\pm 60$ min) |
| Surgery duration | – | $176 \pm 58.2$ min |
| Critical Care Stay | – | 3 (Range 2 - 7) days |
| Hospital Stay | – | 5 (Range 4 - 8) days |

### CABG disrupts normal HPA rhythmicity and elicits a dynamic inflammatory response

All patients exhibited disrupted ACTH and cortisol rhythms following surgery, with hormone concentrations consistently increasing at a time of the day when the control patients’ trend was to decrease. At its maximum (2 hrs post-surgery), mean ACTH levels increased approximately ten-fold compared to physiological levels before decreasing again during the last 6 hrs of sampling. In contrast, mean cortisol increased steadily from the end of surgery, approximately doubling mean physiological levels by the end of sampling (8 hrs post-surgery). The mean levels of inflammatory mediators IL6, TNFα, IL10 and IL8 exhibited dynamic responses as well, but these typically started during the second half of the surgery (Fig. 1D).

In the healthy controls, ACTH and cortisol exhibited regular ultradian pulsatility (T_u_ = 2.5-3 hrs) with a circadian modulated amplitude (Fig. S1). The area under the curve (AUC) for ACTH and cortisol between 8 am and 8 pm (equivalent to the 12-hr sampling time interval for patients) was consistent across controls, with AUC_ACTH_ = 196.65 ± 42.81 ng·hr/L and AUC_CORT_ = 2896.62 ± 221.04 nM·hr. During and after CABG, the dynamic responses of these hormones and inflammatory mediators varied across patients (Fig. S2), with an almost 6-fold increase in total ACTH secretion with respect to controls (AUC_ACTH_ = 1228.82 ± 831.59 ng·hr/L) but only about a 2-fold increase in total cortisol secretion compared to controls (AUC_CORT_ = 6473.7 ± 1181.67 nM·hr). To better characterise the dynamic responses elicited by surgery, we quantified the synchrony between ACTH and cortisol time series by means of non-stationary statistics such as time-dependent hormone ratios and their dynamic range, time-dependent cross-correlations, angle and instantaneous phase synchrony (IPS). These results showed that CABG induces different ACTH/cortisol ratios compared to controls (Figs. S1 and S3), and supports previous work that demonstrates a dynamic dissociation between these hormones in this context^4,9^. The time-dependent Pearson correlation coefficient showed longer-lasting correlated and anti-correlated events between ACTH and cortisol compared to controls, while the angle and IPS showed a loss of phase coherence between ACTH and cortisol rhythms (Figs. S1 and S3). Taken together, these results suggest that the dynamic dissociation between these hormones could be explained by a misalignment of secretion between the pituitary and adrenal glands (due to signals from outside the HPA axis such as inflammatory mediators), an increase in cortisol half-life (due to reduced inactivation into cortisone), or a combination of both.

### Identifying three distinct phenotypes of HPA axis activity following CABG

We investigated the dynamic dissociation between ACTH and cortisol levels by means of cross-correlation algorithms that quantify the synchrony between time series data^14,15^.

Specifically, we calculated: (1) the time-lagged cross correlation (TLCC) to estimate the peak association between ACTH and cortisol; and (2) the rolling window time-lagged cross correlation (RWTLCC) for a time-dependent quantification of the lag between ACTH and cortisol. These algorithms differ from traditional cross-correlations in that they consider the non-stationary nature of variables (see Methods), and allow us to characterise the inter-individual variability of HPA axis responses in patients.

The TLCC and RWTLCC showed that the healthy control group had a high peak association between ACTH and cortisol, with ACTH leading cortisol by a mean lag of *μ*_lag_ = 10 min which remained stable across all epochs (Fig. 2A and Fig. S1). In contrast, we found that the patients experienced different lags and degrees of phase synchrony loss between ACTH and cortisol. Combining the period, TLCC and RWTLCC allowed us to group the dynamic responses of the HPA axis into three categories (Figs. 2B-D) according to the type of dynamic dissociation between ACTH and cortisol elicited by CABG as follows:

- ***Two-pulse group:*** These patients showed two pulses of ACTH and cortisol with longer than physiological periodicity (T_u_ = 5-6 hrs) but preserving a near-physiological peak association (*μ*_lag_ = 13 min) and a stable phase synchrony across the entire 12 hr long sampling (Fig. 2B).
- ***Multiple-pulse group:*** These patients showed multiple pulses of ACTH and cortisol (T_u_ = 2 hrs) with strong peak dissociation (*μ*_lag_ = 106 min) and partial phase synchrony during a portion of the 12-hr long sampling (Fig. 2C).
- ***Single-pulse group:*** These patients showed a single large excursion of ACTH and cortisol with strong peak dissociation (*μ*_lag_ = 77 min) and unstable phase synchrony across the entire 12-hr long sampling (Fig. 2D).

**Figure 2.**
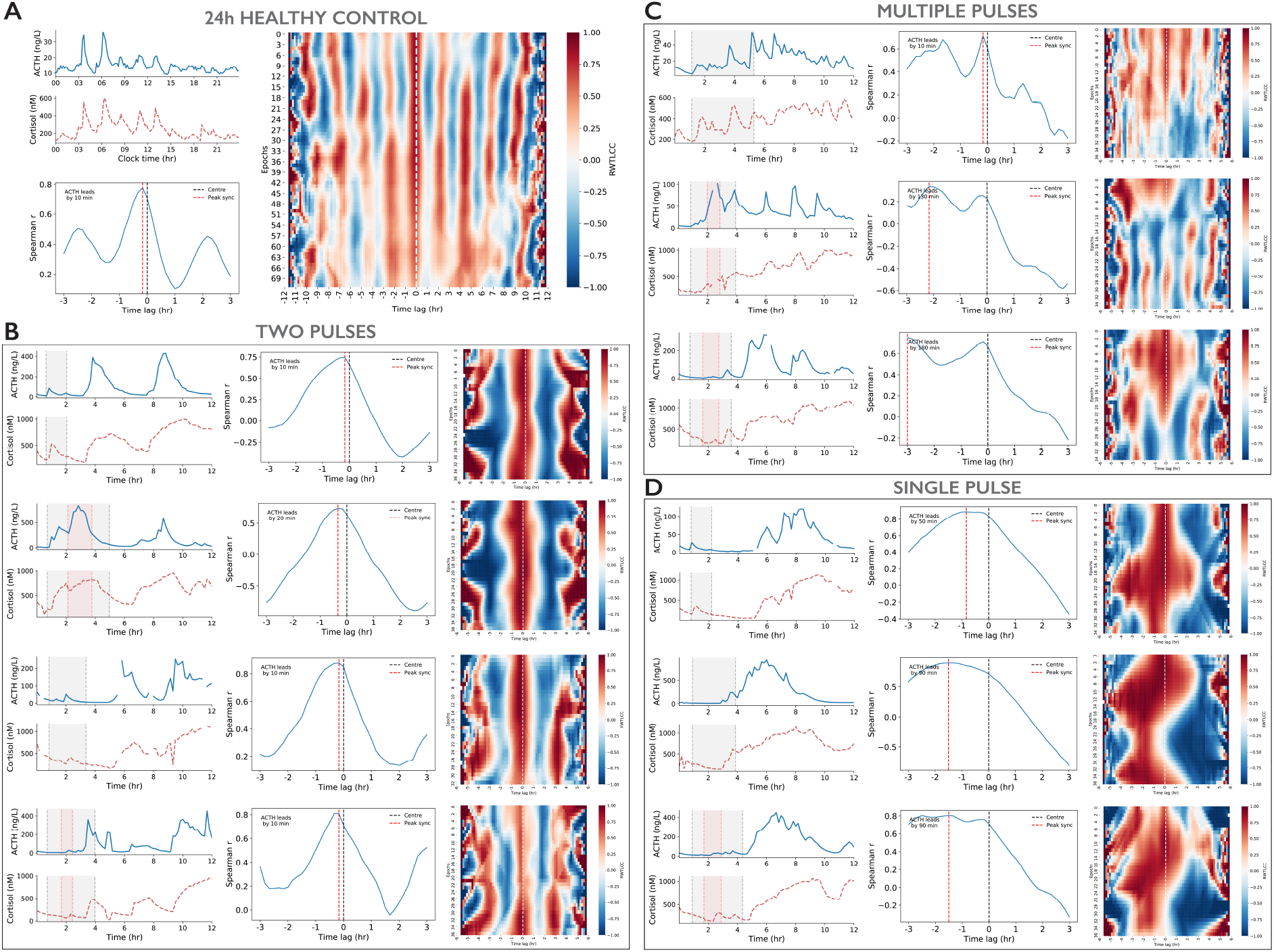
Dynamic responses of ACTH and cortisol during and after CABG (n=10). **A)** Ultradian rhythms with circadian modulated amplitude in one healthy control. While the time-lagged cross correlation (TLCC) quantifies peak association between ACTH and cortisol, the rolling window TLCC heat map (RWTLCC) characterises the phase synchrony across all epochs (elapsed time increases downwards). We use these techniques to identify three types of CABG-induced cortisol profile phenotype: **B) Two-pulses**: characterised by two ACTH and cortisol pulses of supra-physiological amplitude and period, but without loss of phase synchrony, **C) Multiple-pulses**: characterised by multiple ACTH and cortisol pulses of increased amplitude but near normal periodicity, with loss of peak association but no apparent loss of phase synchrony, and **D) Single-pulse**: characterised by a single ACTH and cortisol pulse of supra-physiological amplitude, with loss of both peak association and phase synchrony. Grey (red) shaded areas in the hormone time series indicate duration of CABG surgery (CPB).

These techniques allow us to describe the inter-individual variability of dynamic responses to CABG in quantitative terms. They suggest that patients may fall into different categories depending on the type and degree of the dynamic dissociation between ACTH and cortisol following surgery. Namely, the two-pulse group experiences lesser dynamic dissociation, the single-pulse group experiences greater dynamic dissociation, and the multiple-pulses group falls in the middle. These statistical methods do not reveal what physiological processes underpin this variability. They also do not show whether they originate from the interplay between the HPA axis and inflammatory responses. To provide insight into how different physiological processes may contribute to the altered profiles of cortisol, we developed a mathematical model of cortisol concentration.

### A model of cortisol concentration accounts for the dynamic changes of the HPA axis following CABG

The model assumes a delayed (10 minutes) ACTH input to a hypothetical adrenal gland underpinning cortisol production (implicitly including peripheral conversion from cortisone), cortisol turnover, and the adrenal sensitivity to ACTH stimulation. It also assumes a two-compartment, open-loop architecture where each compartment accounts for fast and slow processes contributing to cortisol dynamics, respectively^16^. In the model (Fig. 3A and Methods), parameters *p_f_* and *p_s_* account for fast and slow cortisol production respectively (including adrenal maximum secretory capacity and peripheral conversion from cortisone); *1/λ_f_* and *1/λ_s_* are the fast and slow cortisol turnover rates, respectively; *K_A_* is the adrenal sensitivity to ACTH stimulation; and *m* is the Hill coefficient denoting the steepness of the sigmoidal function used to represent the adrenal response to ACTH.

**Figure 3.**
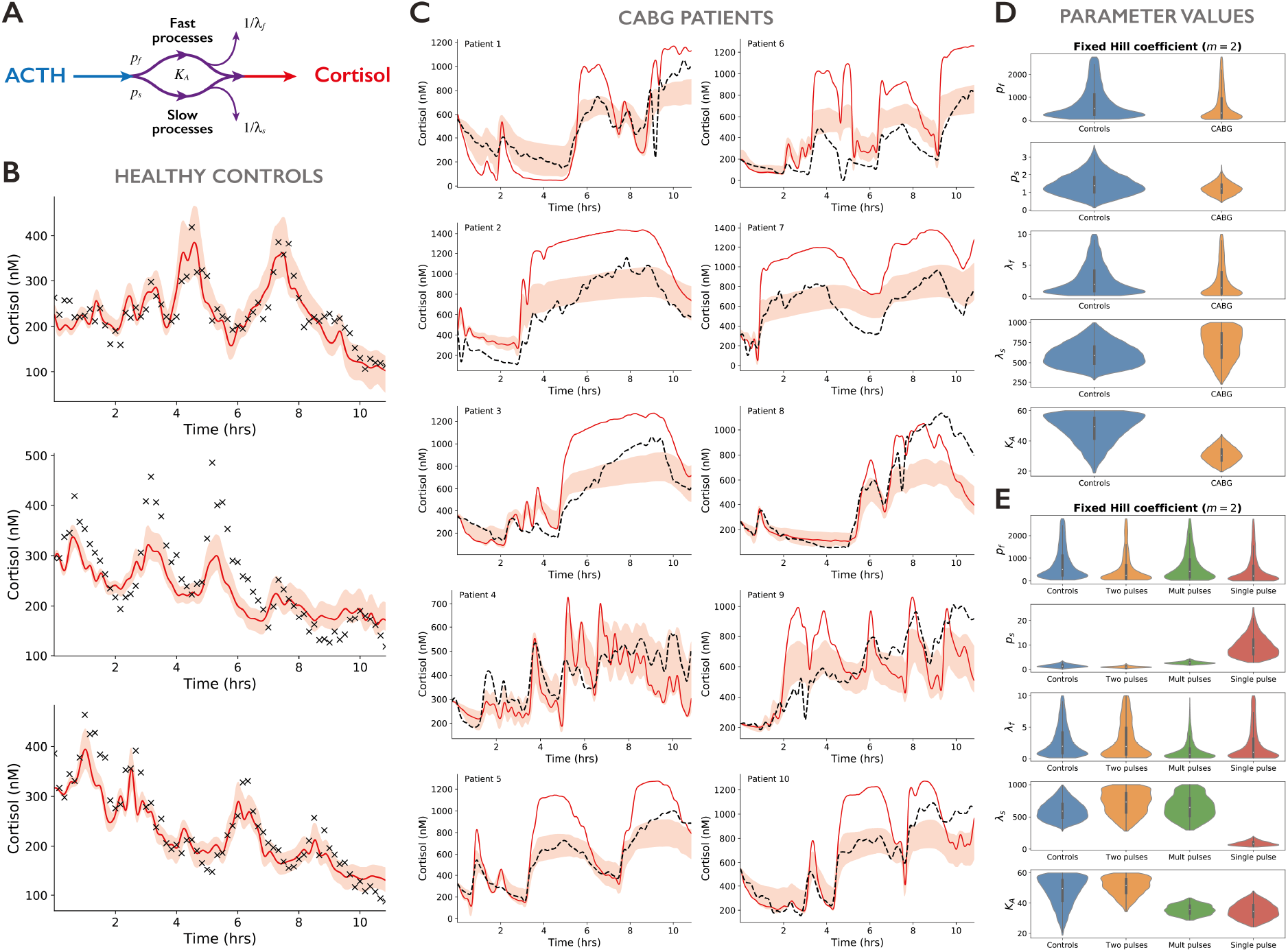
Mathematical model of cortisol activity. **A)** Schematic illustrating the mathematical model. About 11 hrs of ACTH data was used as input into fast and slow cortisol activity compartments. The model was calibrated against **B)** healthy controls, and its predictions compared against **C)** patient data. Black dots and broken lines are cortisol data, shaded red areas are model predictions, and red continuous lines are the predictions from the median of parameter values. **D)** Distributions of best parameter fits with fixed Hill coefficient (m=2) for controls, all patients grouped together, and patients grouped by cortisol profile pattern type.

We performed computer simulations of the model and calibrated it to physiological data. To do this, we used the ACTH data as an input and assessed the ability of the model to accurately capture the cortisol output trajectories over 1 million sets of parameter values. The motivation for this was two-fold. First, we aimed to determine whether parameters that fit the control cortisol trajectories could also explain the cortisol trajectories in patients undergoing CABG without the cytokine input. If this were the case, then we could assume that any potential influence of inflammatory cytokines would occur at the level of the pituitary rather than the adrenal glands (Fig. 1B). Second, if the parameters from the controls were not a good fit to the data from patients having CABG, then we could assess the changes required to achieve a good fit. This would help identify potential targets where cytokines could be influencing cortisol activity, for example by regulating secretion from the adrenal glands or activity in peripheral tissues (Fig. 1C).

#### Modelling controls and CABG patients

To calibrate the model, over 1 million sets of candidate parameter values were randomly selected within a latin hypercube and tested to fit the control data. The cost function (see Methods) used during the fitting process was evaluated for each of the controls and each of the CABG patients. Table S1 includes the ranges of tested values for each model parameter and the distribution of optimal parameter values that fit the control group. In order to establish a baseline set of parameters, we first identified the best fit parameter set (single lowest cost function evaluation) as well as the group of parameter sets whose cost function evaluation was within 10% of the lowest evaluation. We found that most of the fits to controls corresponded to scenarios where the Hill coefficient was fixed to *m* = 2; therefore, we used this value for the rest of the analysis. We also found that the error was low for all controls (median error 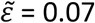 across the fit parameter values) (Fig. 3B), showing that the model accurately captures the control data. Having many parameter sets that fit the control data with low error implies that there are many parameter sets that can fit all controls simultaneously. The dark red line in Fig. 3B shows the cortisol trajectories predicted by the model when using the median values of the distribution of parameters that best fit all controls 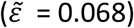.

The median error for the fits to the CABG patient data was 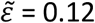, nearly double that of the fits to the controls. Fig. 3C shows the predicted cortisol trajectories using these fits against the CABG patient data. This highlights that while these trajectories were not as good as the fits to the controls, they still follow the cortisol trend observed in CABG patients. The predicted trajectories using the fits to the patient data perform better than when using the median parameter values fitted to the controls (Table S1) to simulate the CABG cases (dark red line Fig 3B, 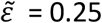). In other words, when the model parameters are fitted to the CABG patient group, the predicted trajectories are closer to the patient data than when using the median parameter values obtained from fitting the model to the control group. To better characterise the differences between the control and patient groups, we used violin plots to represent the distributions of optimal parameter values resulting from fitting the model to data (Fig. 3D). The cortisol production rates *p_f_* and *p_s_*, as well as the fast cortisol half-life *λ_f_*, had similar statistical moments and shape of their distributions. In contrast, while the slow cortisol half-life *λ_s_* was found to have similar statistical moments, its distribution was different between controls and patients, leaning towards higher values (implying slower cortisol turnover) in CABG patients. A more striking difference was observed in the adrenal sensitivity *K_A_*, where the estimated distribution had a lower, narrower distribution in the CABG case compared to controls. Taken together, this suggests that the same model explaining the cortisol dynamics observed in controls can also explain that in patients having CABG by assuming a slower cortisol turnover and an increased adrenal sensitivity to ACTH stimulation.

#### Modelling CABG patients with different phenotypes of cortisol profile

To investigate the origin of the distinct dynamic dissociation between ACTH and cortisol observed across the three groups of patients (two-pulses, multiple-pulses, and single-pulse), we fitted the model to identify optimal parameter values for each group (Table S2). The fits were marginally improved compared to the fits to all CABG patients pooled together, with the error decreasing to 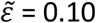 for the two-pulses group, 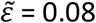 for the multiple-pulses group, and 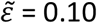 for the single-pulse group (Table 2). The shapes of distributions of parameter values associated with fast processes (*p_f_*, *λ_f_*) were also very similar across all CABG groups and between these and the control group (Fig. 3E). For parameters associated with slow processes, the cortisol production rate *p_s_* and half-life *λ_s_* were predicted to remain within the same dynamic range (although slightly higher) as controls for the two-pulses and multiple-pulses CABG groups (Table S2). The shape of the *λ_s_* distribution tending towards higher values in these patient groups. In contrast, the single-pulse CABG group was predicted to have a higher cortisol production rate *p_s_*, but a lower half-life *λ_s_*. The adrenal sensitivity *K_A_* was similar between the control and the two-pulses CABG group, but was predicted to have lower values (implying higher adrenal sensitivity) for the single-pulse and multiple-pulses groups (Fig. 3E). Taken together, these results suggest that the degree of cortisol pattern change observed across the different CABG groups could originate from a combination of disrupted physiological processes. Specifically, the estimated distributions of parameter values for the two-pulses group maintain similar statistical moments to the healthy controls, only changing the shape of the *λ_s_* distribution. The distributions for the multiple-pulses group are similar to the two-pulses group, except that the model also predicts an increased adrenal sensitivity (lower *K_A_*). Finally, the distributions for the single-pulse group show that not only is the adrenal sensitivity increased, but also the cortisol secretion *p_s_* and its turnover rate 1/*λ_s_*. The dynamic range and shape of the distribution of *K_A_* predicted for the two-pulses CABG group was very close to controls, suggesting that the adrenal sensitivity to ACTH is the same for these patients.

**Table 2.** Median error values of the parameter sets fit to the controls, patient group (CABG), and each subgroup.

| Parameter | Median Error of Cost Function Evaluations |  |  |  |  |
| --- | --- | --- | --- | --- | --- |
|  | Control | CABG | Two-Pulses | Multiple-Pulses | Single-Pulse |
| $m = 2$ | 0.07 | 0.12 | 0.10 | 0.08 | 0.10 |
| $m = 2, K_A = 50.84$ | 0.07 | 0.14 | - | - | - |

#### Model predictions under fixed adrenal sensitivity

To further investigate the origin of the dynamic dissociation between ACTH and cortisol, we used the assumption that CABG surgery does not markedly change the adrenal sensitivity to ACTH stimuli. This implies that such changes are unlikely to take place at the time scale of hours after surgery but may result from long term critical illness^17,18^. To do this, we fixed the adrenal sensitivity parameter to the median value estimated for controls (*K_A_* = 50.28) and identified the distributions of parameter values as before (Table S3). The model predicted low error trajectories for both the controls 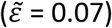 and the CABG group 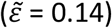. This marginally changed with respect to the baseline parameter fitting (compare Figs. 4A and 4B with Figs. 3B and 3C). The optimal parameter sets, when compared between the control and CABG group, showed these to be strikingly similar. The exception was the shape of the distribution for the half-life parameter *λ_s_*, which tended towards higher values (Fig. 4C). This suggests that, if adrenal sensitivity does not change after CABG, the key physiological process underpinning dynamic dissociation in patients would likely be a slower cortisol turnover rate^19^.

**Figure 4.**
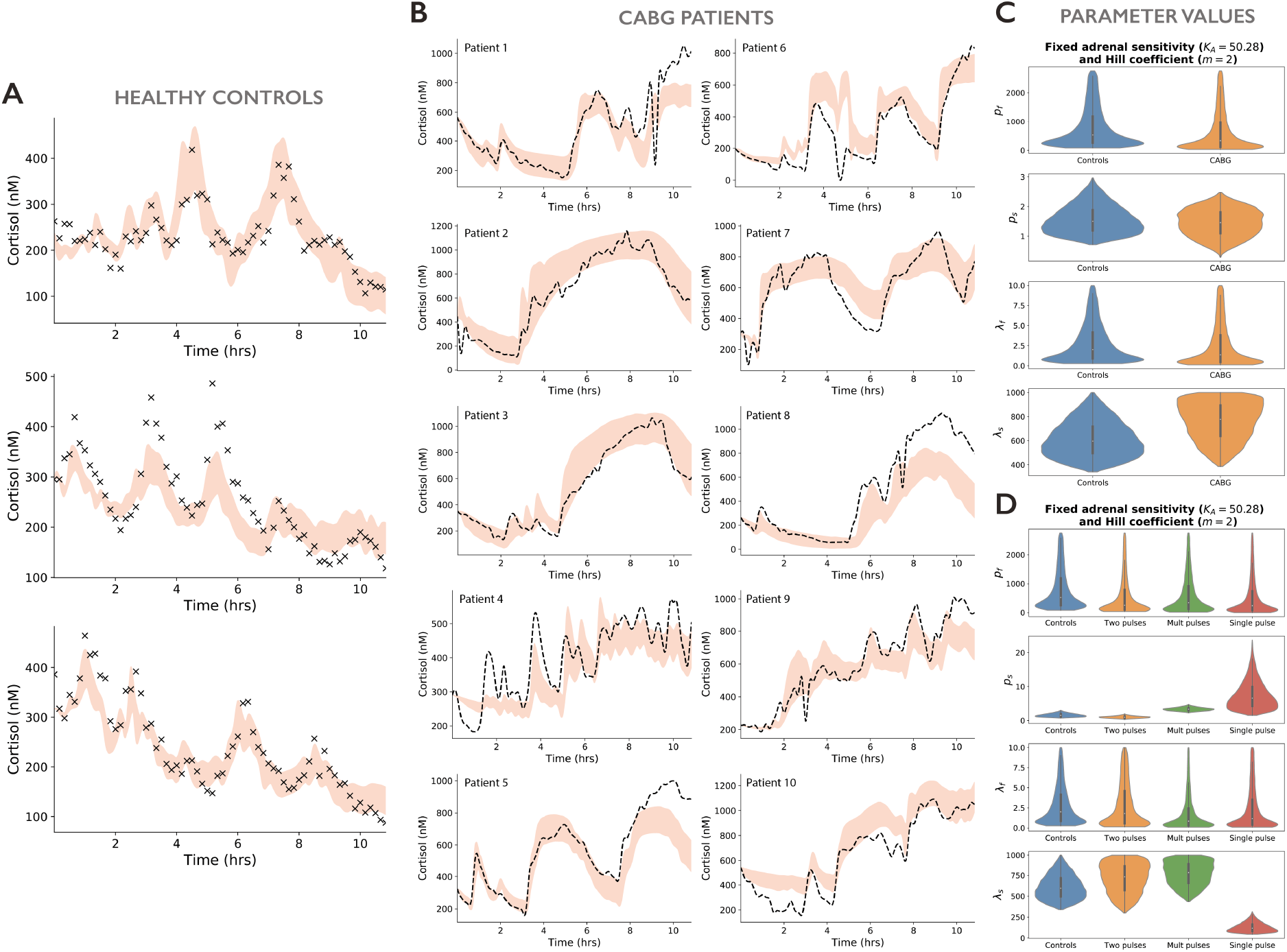
Model re-calibration assuming fixed adrenal sensitivity and Hill coefficient. Model predictions after re-calibration for **A)** healthy controls, and **B)** patient data. Black dots and broken lines are cortisol data, shaded red areas are model predictions. **C-D)** Distributions of best parameter fits with fixed adrenal sensitivity (K_A_=50.28) and fixed Hill coefficient (m=2) for controls, all patients grouped together, and patients grouped by their type of chronodisruption.

Next, we investigated the model predictions for each CABG group under the assumption of fixed adrenal sensitivity. As before, the shapes of distributions for optimal parameter values associated with fast processes (*p_f_*, *λ_f_*) were very similar across all CABG groups and the control group (Fig. 4D). For parameters associated with slow processes, the cortisol production rate *p_s_* and half-life *λ_s_* were predicted to remain within the same dynamic range (albeit slightly higher) as controls for the two-pulses and multiple-pulses CABG groups (Table S3), with the shape of the *λ_s_* distribution leaning towards higher values in these patient groups. Consistent with the variable *K_A_* scenario, the single-pulse CABG group was again predicted to have a higher cortisol production rate *p_s_*, but a shorter half-life *λ_s_*.

### Inflammatory mediators may underpin the dynamic changes in the interaction of ACTH and cortisol following CABG

While the model can predict cortisol dynamics driven by ACTH inputs in both healthy controls and CABG patients, minor discrepancies between the model predictions and data persist. We now explore whether these discrepancies may be related to the effect of inflammatory mediators, which have not been explicitly included in the mathematical model to this point. To do this, we first performed a principal component analysis (PCA) on the trajectories of inflammatory mediators following CABG (Fig. S2). The PCA identified IL6 and TNFα as the cytokines containing the most information about the inflammatory response (Fig. S4). IL6 and TNFα have previously been shown to be involved in the regulation of 11β-HSD1/2 enzymatic cortisol to cortisone inter-conversion^20–22^ (Fig. 1C).

We then calculated the time-varying residual error between the model predictions and the cortisol trajectories. Following Z-score normalisation, we extracted correlations between the model error and the inflammatory mediator dynamics. This allowed us to compare each cytokine time-varying trajectory with the error of predicted cortisol levels for each patient, and visually inspect where the model discrepancies with data could possibly be explained by cytokine dynamics (Fig. 5A). Since inflammatory mediators are likely to act together in regulating the HPA axis, we calculated correlations not only for cytokines IL6 and TNFα identified by the PCA, but also for IL8, IL10, and additive combinations of them. This is summarised in Fig. 5B, where each patient’s correlation between their predicted cortisol trajectory error and inflammatory mediators are represented as a row in a correlation matrix heat map. Patients 2, 3 and 8 – belonging to the single-pulse group that experienced the greatest chronodisruption – showed some of the highest positive (patients 2 and 3) and negative (patient 8) correlation coefficients. In contrast, patients 1, 5, 6 and 7 belonging to the two-pulses group showed positive but weak correlations with inflammatory mediators, while the remaining patients belonging to the multiple-pulses group showed a mix of low positive, low negative and near null correlations. These results suggest that the greater dynamic dissociation of ACTH and cortisol observed in the single-pulse group may be explained by cytokine-induced changes in the cortisol production rate and half-life. In other words, the correlations between the model residual error and cytokines can help identify the patients in which the inflammatory response is more likely to underpin the dynamic dissociation between ACTH and cortisol. This coincides with the highest correlations observed in groups of patients that also exhibit greater disruptions in HPA axis activity following CABG surgery.

**Figure 5.**
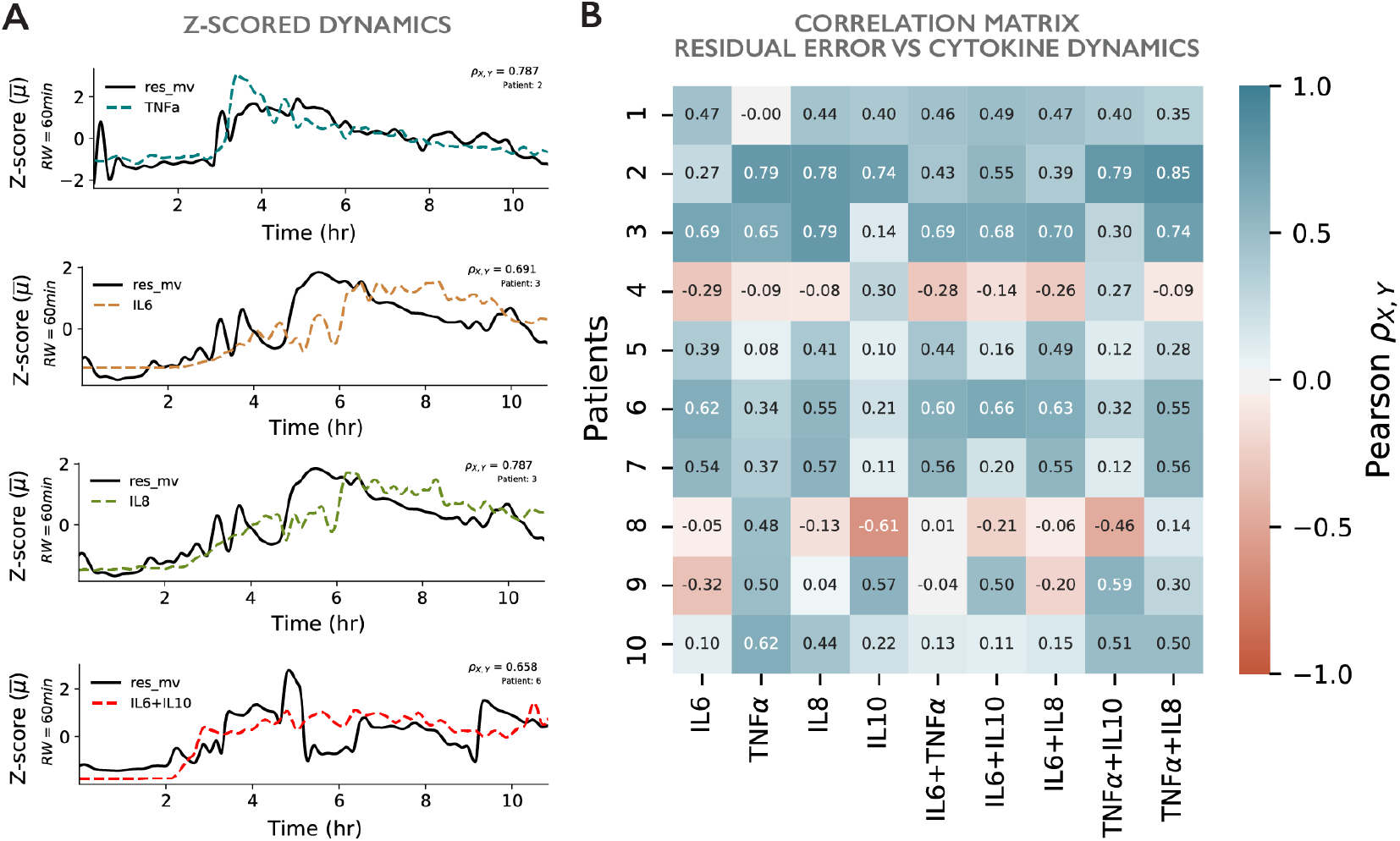
Correlation between model predictions error and cytokine dynamics. **A)** Examples of Z-scored dynamics of the model residual error (predictions minus data) and selected inflammatory mediators for three patients. **B)** Correlation matrix between the model residual error and the dynamics of inflammatory mediators for each patient. Patients in two-pulses group: 1, 5, 6 and 7. Patients in multiple-pulses group: 4, 9 and 10. Patients in single-pulse group: 2, 3 and 8.

## DISCUSSION

A maladaptive response to acute systemic inflammation underlies most of the morbidity and mortality from major surgery^23^ and critical illness (most notably sepsis^24^ and more recently COVID-19^25^). Therefore, understanding the mechanisms that modulate this is key to designing clinical diagnostics and interventions that reduce the risk of patient death and morbidity. The HPA axis is one such system that is thought to overall counterbalance inflammation – although this has been difficult to elucidate due to the multiple levels of interaction and sites of feedback between the two systems. High-frequency blood sampling techniques and mathematical models have advanced our understanding of the interaction between the two systems – but previously not in a real-world environment^9,26^. Using a combination of high-frequency blood sampling and statistical analysis, this study has shown that at least three distinct phenotypes of cortisol dynamic profiles emerge after cardiac surgery; single-pulse, two-pulses and multiple-pulses. The mechanisms underpinning these three different phenotypes were inferred using mathematical modelling, with parameter estimations used to predict the contribution of distinct cortisol control mechanisms in patients. Lastly, we identified that inflammatory mediators may only be important in specific patterns of HPA axis activity, paving the way to identify those most at risk of serious harm from inflammation.

We postulated a two-compartment mathematical model of cortisol activity that discriminates between fast and slow processes, with model parameters corresponding to the adrenal sensitivity to ACTH, cortisol production (a contribution of adrenal secretory capacity and peripheral conversion from cortisone), and cortisol turnover and metabolic rates (arising from degradation and inactivation into cortisone). We determined the models with the best fits by assuming a similar adrenal sensitivity between patients undergoing CABG and controls. During fitting, only the parameters associated with fast processes remained invariant between the control and patient groups. This suggests that CABG may not particularly affect cortisol regulatory processes associated with short time scales (e.g. rapid cortisol synthesis and distribution out of the central (plasma) compartment), but only affects slower processes such as enzymatic degradation. When examining the slow parameters, the model predicted that patients in the single-pulse group had a greater slow cortisol production rate and higher slow turnover rate compared to controls and other patient groups. The single-pulse group also had the highest dynamic dissociation between ACTH and cortisol (strong peak dissociation and unstable phase synchrony). The model predicted that patients in the two-pulses group had a similar cortisol production rate and an increased slow turnover rate compared to controls. This group also had the lowest dynamic dissociation between ACTH and cortisol following surgery. Lastly, the model predicted that patients in the multiple-pulses group had an increased slow cortisol production rate and an increased slow turnover rate when compared to controls. This group also had a dynamic dissociation between ACTH and cortisol falling in the middle of the other two phenotypes. Taken together, these results suggest which mechanisms may underpin the three different phenotypes of cortisol activity observed in patients undergoing CABG. Our model also suggests that inflammatory mediators do not play a significant role in regulating circulating levels of cortisol in *all* patients having CABG within the 12 hrs following surgery, as previously postulated^19^. However, in some cases the effect of inflammatory mediators may cause significant disruption to HPA axis dynamics, leading to a large sustained pulse of cortisol.

There are several limitations to consider when interpreting the results of the mathematical model. While the fast and slow time scale separation in the model has been described previously^16,26^, the optimisation is agnostic with respect to the physical mechanisms underlying this separation and only attempts to minimise the cost function. Therefore, some of the fast mechanisms are captured by the slow variable of the model and vice-versa, but this is likely to be minimal. On the other hand, we consider ranges of parameter sets that fit the data well, rather than only considering the single best fit parameter set. This is to minimise concerns of overfitting the data. This means that we can only assess the difference between groups, but not assume that a single group in isolation provides specific estimates of parameter values. Our sample size of patients and healthy participants was small, so we cannot be sure that we have captured the full range of possible HPA axis response patterns that occur after cardiac surgery. It also means we were unable to fully investigate the patient and operative factors that may cause or are associated with the different patterns of physiological processes. We envisage that novel blood-free biosampling technologies^27^ will facilitate sampling a larger number of patients. This will allow a better characterisation of CABG phenotypes and investigate the effects of cytokines over a longer timeframe.

Our model does not consider inflammatory cytokines either as state variables or model inputs (as was the case with ACTH). Nor does it consider any explicit modulation by inflammatory mediators on ACTH – our open loop model architecture assumes such modulation is already accounted for in the resulting ACTH dynamics. Instead, we try to capture potential time-varying modulations of cortisol using parameter changes that remain fixed over time. This could explain the increased error in the CABG patient fits relative to the control fits. This is to be expected given the inflammatory response is only present in CABG patients. However, including inflammatory cytokines as model inputs would not only require additional parameters and therefore increase the risk of overfitting, but would also require detailed knowledge of the interactions between the HPA axis and inflammatory pathways. Although some advances have been made in modelling these interactions^9,28^, uncovering the network architecture between these pathways after major surgery and in critical illness has so far not been achieved. To do so will require a larger cohort of patients and a combination of mathematical modelling and machine learning techniques^29^.

We have been able to show that that there is not a simple graded HPA response to cardiac surgery. There are major dynamic changes involving not only the concentration of ACTH and cortisol but also their pattern of secretion. Using novel mathematical techniques, we show that CABG patients exhibit three different patterns of HPA axis response, which reflect different underlying physiological changes in adrenal sensitivity, cortisol production and turnover. Inflammatory mediators appear to be driving changes in only one of these patterns – the pattern of a single-pulse implying loss of ultradian pulsatility – despite being postulated as the likely cause for the elevated cortisol seen in all types of systemic inflammation^19,28^. We suggest that the different patterns we have observed may represent different “severities” of response to the surgery. An adequately powered study is now required to investigate whether and how these patterns are correlated with clinical outcomes. This will be critical to establish whether the patterns could be used for risk stratification after surgery. This study also shows that the existing model of corticosteroid physiology used for diagnosis and prognosis after major surgery and in critical illness^30^ may only represent mean values within a population rather than the responses of individuals or groups of individuals. Improved diagnostics based on individual or subgroup responses is likely to lead to greater precision in diagnosis and more targeted interventions.

## METHODS

The study was reviewed and approved by the UK National Research Ethics Service (NRES, Ref: 11/H0107/9) and Health Research Authority (HRA). Female sex hormone cycles are known to affect the HPA-axis^31,32^ and for this reason, female participants were excluded.

### Subjects – patients

Ten patients were recruited prior to surgery from a single cardiac surgical centre and met the following inclusion criteria: male, aged 18 – 80 years undergoing first time, elective CABG carried out via median sternotomy. Patients were excluded if they met any of the following exclusion criteria: emergency operation, previous sternotomy, myocardial infarction within the last month, concomitant procedure with CABG, left ventricular ejection fraction <30%, operation to be carried out by other incision than median sternotomy (e.g. left thoracotomy), use of exogenous corticosteroids (including inhalers), history of adrenal or pituitary disease.

### Subjects – Healthy controls

Healthy controls were taken from previously published studies on healthy subjects^26,33^. In brief, healthy males were recruited with informed consent. Participants were excluded if they had recent (<10 days) trans-meridian travel and if they were exposed to exogenous glucocorticoids in the previous 3 months. Participants maintained conventional work and sleeping patterns and were sampled from peripheral veins in a clinical study unit. Meals were provided at 0800, 1230, and 1730 hrs, and room lights were turned off between 2200 and 2400 hrs depending upon individual sleeping habits.

### Collection and processing of blood samples

All surgical procedures were scheduled first of the day (beginning at 8am). Surgical patients had blood sampled for 12 hours via their *insitu* vascular catheters from first cannulation using a needle-free, closed-loop sampling system (Edwards VAMP. Edwards Life Sciences Corp, Irvine, CA. USA). Healthy controls were sampled from peripheral veins by either hand sampling or an automated blood sampling system^33^. Total serum cortisol, ACTH and four inflammatory mediators (IL1α, IL2, IL4, IL6, IL8, IL10 and TNFα) were sampled at 10-minute intervals. Cortisol samples were collected in BD vacutainer SST Advance tubes (Becton, Dickinson and Company, Oxford. UK) and were processed immediately after centrifugation. Samples for ACTH were collected in chilled 2ml EDTA tubes and stored on ice for a maximum of 60 minutes before centrifugation at 4°C and then stored at −80°C until assay. Total cortisol and ACTH were measured by solid phase, chemo-luminescent enzyme linked immunoassay (ECLIA) using the Cobas e602 modular analyser (Roche Diagnostics Ltd, West Sussex, UK). Measuring limits for the cortisol assay were 0.5 – 1750nmol/L (intra- and inter-assay coefficients of variation (COV): 1.5 – 1.7% and 1.8 – 2.8% respectively) and for the ACTH assay were 1.0 – 2000 pg/ml (intra- and interassay COV: 0.6 – 2.7% and 3.5 – 5.4% respectively). Inflammatory mediators were collected in the same BD vacutainer SST advance tubes as cortisol. After centrifugation, aliquots were stored at −80°C until assay. Inflammatory mediators were assayed using the Luminex® Multiplex system (ThermoFisher Scientific - Waltham. MA)

### Statistical analysis

The time-lagged cross correlation (TLCC) was used to quantify the strength of peak association (Spearman r) and to estimate the time lag between ACTH and cortisol dynamics. The rolling window time-lagged cross correlation (RWTLCC) was used to quantify the time lag when the peak association occurred for a range of elapsed time values from the beginning of the time series. While the TLCC can be regarded as a global metric that focuses on ACTH and cortisol peaks to estimate the overall time lag between these hormones, the RWTLCC splits the time series into epochs, thus allowing a time-dependent quantification of the lag between ACTH and cortisol. The RWTLCC can be better visualised through a heat map, where vertical stripes emerge naturally when the association between variables becomes stable over time and therefore their association is stationary. This is the case of healthy controls where ACTH and cortisol show a strong, stable phase synchrony (Fig. 2A).

Simpler, stationary cross-correlation methods were used to estimate the potential effects of inflammatory mediators not accounted for in the model. These involved a Z-score normalisation of the residual error of the model predictions vs cortisol data, and calculating its cross-correlation with the Z-score normalisation of the inflammatory mediators (Fig. 5A). The results are summarised in a cross-correlation matrix for each patient, each cytokine, and some key combinations of these cytokines (Fig. 5B). We also performed a pairwise principal component analysis of the hormones ACTH and cortisol, as well as the inflammatory cytokines, to determine how similar their dynamic profiles were to one another. We did this on Z-scored data and used the MATLAB function pca to identify the variance explained by the two principal components, which were identified as IL6 and TNFα. For each pair, we found the principal components by concatenating all patient data together. Therefore, the percentages of variance were calculated at the patient group level (Fig. S4).

### Mathematical model

We use a two-compartment mathematical model for cortisol, that depends on ACTH. Here, the ACTH data are treated as input and the model output is cortisol. We then use an optimisation algorithm to identify suitable parameter values of the model for an individual. The model is represented by the equation:

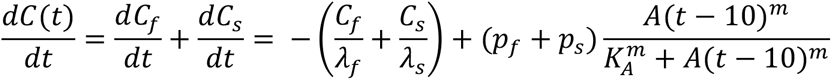

The model splits the cortisol dynamics *C(t)* as the contribution of two compartments, *C_f_* and *C_s_*, respectively accounting for fast and slow time scales for cortisol dynamics. Parameters *p_f_* and *p_s_* respectively account for the fast and slow cortisol production rates (including adrenal maximum secretory capacity and conversion from cortisone); 1/*λ_f_* and 1/*λ_s_* are the fast and slow cortisol turnover rates, respectively; 1/*K_A_* is the adrenal sensitivity to ACTH stimulation; and *m* is an integer denoting the power of the Hill function. *A* is the ACTH data, for which we assume a 10-minute delay before its effects are reflected onto cortisol^9^ (Fig. 2A). Overall, the model has six parameters, some of which we fix at different stages in the study. The initial condition for the model is chosen by setting the fast compartment *C_f_* to quasi-equilibrium, then *C_s_*(0) = max (*C_data_*(0) – *C_f_*(0),0).

### Optimisation

We use the cost function:

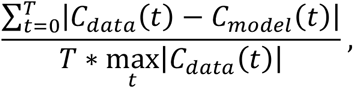

where *C_data_* is the cortisol data for a given control or patient and *C_model_* is the model cortisol output for a given parameter set (using the control or patient ACTH data as input). We derive the sample parameter sets from a latin hypercube of 1024^2^ points, capture the cost function for each patient or control and for each parameter set. This results in two matrices:

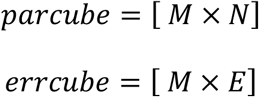

Where *M* = 1024^2^ the number of parameter sets on the hypercube, *N* = 6 the number of parameters, and *E* = 13 the number of patients (10) plus controls (3). *Parcube* is the list of parameters and *errcube* is the list of errors. We can then sort *errcube* and *parcube* by an individual control or patient to find the best-fit parameter choices. In some cases, we also sort by the patient group as a whole (or a subgroup of patients) or the control group. In this case, we use the mean-square of the error (MSE) across the given group to sort the parameters. This allows us to identify parameter sets that are best-fit for groups of multiple individuals. In some cases, we also fix some of the parameters and run a new latin hypercube. For example, we identified the best choice of the exponent parameter *m* using an initial hypercube. We found that most good fits were for *m* = 2, so we fixed that parameter and identified a new hypercube of 5 parameters for subsequent simulations. We identify good fits as any solution with an error value less than 10% greater than the best-fit error value. Therefore, we identify a distribution of good-fit solutions to minimize issues related to overfitting.

## DATA AVAILABILITY

Raw data on hormone profiles, inflammatory mediators and surgery times, together with the time series analysis implemented in Python 3.6 for CABG phenotyping is available at https://gitlab.com/ezavala1/CABG_phenotypes

The full model and optimisation pipeline was implemented in MATLAB and is available at https://github.com/dgalvis/heart_surgery_model

## ACKNOWLEDGMENTS

This study was funded by the British Heart Foundation, the Engineering and Physical Sciences Research Council (EPSRC), a Medical Research Council (MRC) fellowship (MR/P014747/1) to E.Z., a Medical Research Council (MRC) fellowship (MR/N008936/1) to J.J.W., the NIHR Biomedical Research Centre at University Hospitals Bristol and Weston NHS Foundation Trust and the University of Bristol. The views expressed are those of the author(s) and not necessarily those of the NIHR or the Department of Health and Social Care.

## AUTHOR CONTRIBUTIONS

DG, EZ, JJW, SLL and BG conceived and designed the study. BG, JE, CAR and KP collected the data. DG, EZ, JJW and TU developed the statistical analysis and mathematical model. DG and EZ performed the analytical calculations and computer simulations. DG, EZ, JJW and BG wrote and edited the manuscript. SLL, GDA, JE, CAR and KP edited the manuscript.

## COMPETING INTERESTS

The authors declare no competing interests.

## Notes

https://gitlab.com/ezavala1/CABG_phenotypes

https://github.com/dgalvis/heart_surgery_model

### Competing Interest Statement

The authors have declared no competing interest.

