## Supplementary Figs and Tables for "The dynamic interaction of systemic inflammation and the hypothalamic-pituitary-adrenal (HPA) axis during and after major surgery"

### Supplementary Figures

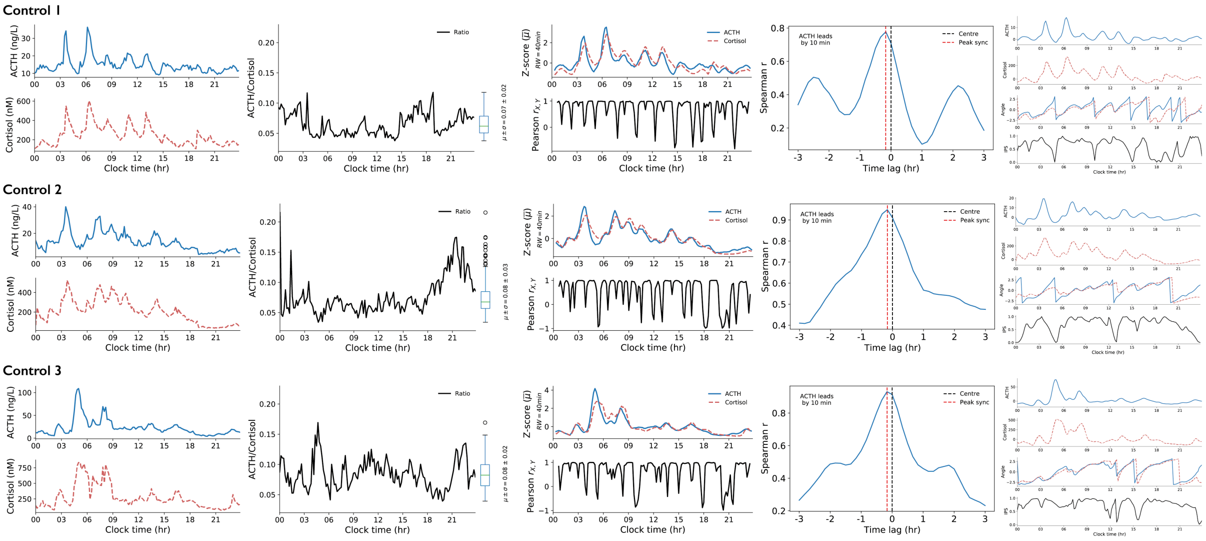

**Figure S1. HPA axis activity of healthy controls.** From left to right: 24hr plasma ACTH and cortisol profiles from 3 healthy individuals, time-varying ACTH to cortisol ratios and their dynamic range, z-score normalisations and their time-varying Pearson correlation, ACTH to cortisol lag as revealed by the time-lagged cross-correlation (TLCC), and time-varying angle and instantaneous phase synchrony (IPS) between ACTH and cortisol.

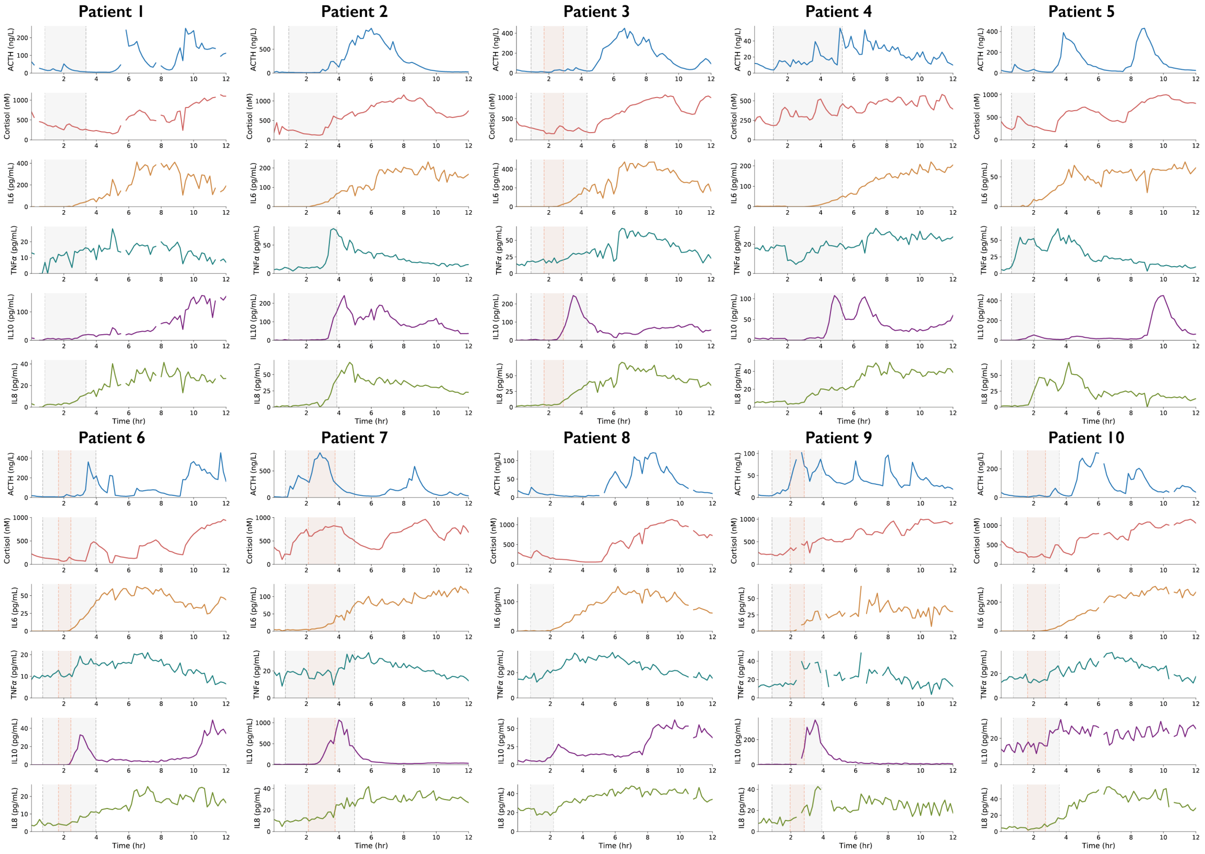

**Figure S2. HPA axis activity and inflammatory response of CABG patients.** 12hr time course of ACTH, cortisol, IL6, TNF$\alpha$, IL10 and IL8, during and after CABG. The elapsed time of surgery (CPB) is shown as a shaded grey (red) area.

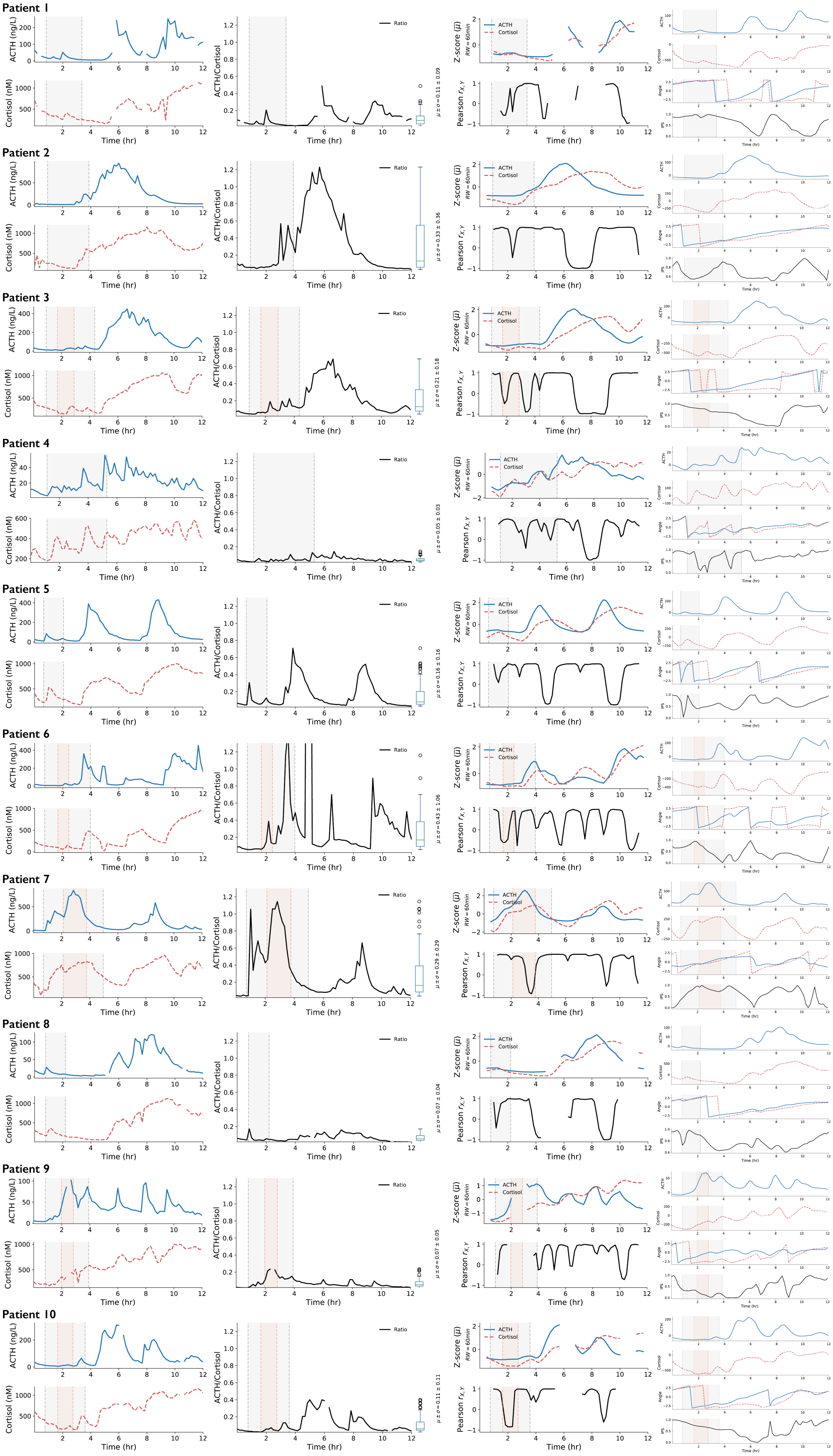

Patient 1
Patient 2
Patient 3
Patient 4
Patient 5
Patient 6
Patient 7
Patient 8
Patient 9
Patient 10
ACTH (ng/L)
Cortisol (nM)
Time (hr)
ACTH/Cortisol
Ratio
Z-score ($\bar{\mu}$)
RW = 60min
ACTH
Cortisol
Pearson $r_{X,Y}$
Angle
IPS
$\mu \pm \sigma = 0.11 \pm 0.09$
$\mu \pm \sigma = 0.33 \pm 0.36$
$\mu \pm \sigma = 0.21 \pm 0.18$
$\mu \pm \sigma = 0.05 \pm 0.03$
$\mu \pm \sigma = 0.16 \pm 0.16$
$\mu \pm \sigma = 0.43 \pm 1.06$
$\mu \pm \sigma = 0.29 \pm 0.29$
$\mu \pm \sigma = 0.07 \pm 0.04$
$\mu \pm \sigma = 0.07 \pm 0.05$
$\mu \pm \sigma = 0.11 \pm 0.11$

**Figure S3. HPA axis activity of CABG patients.** From left to right: 12hr plasma ACTH and cortisol profiles from 10 patients, time-varying ACTH to cortisol ratios and their dynamic range, z-score normalisations and their time-varying Pearson correlation, and time-varying angle and instantaneous phase synchrony (IPS) between ACTH and cortisol. The elapsed time of surgery (CPB) is shown as a shaded grey (red) area.

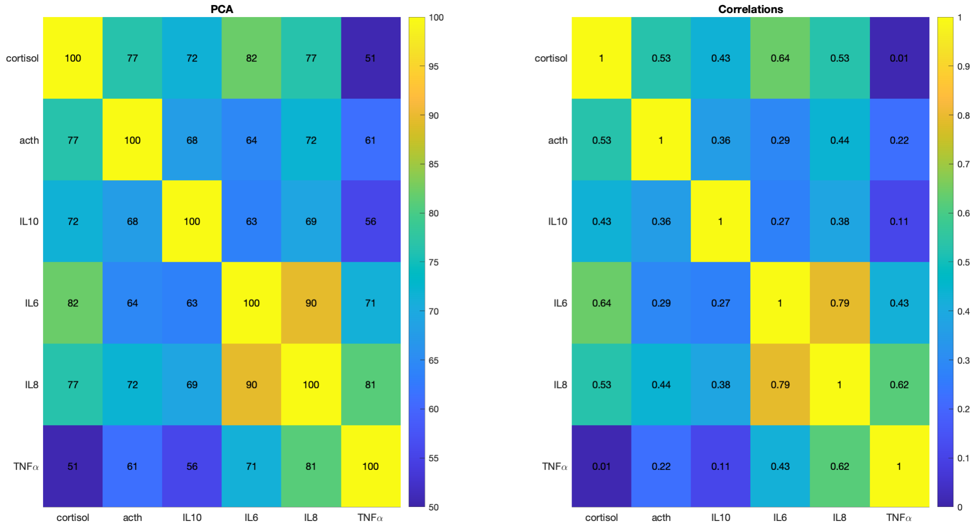


**Figure S4. Principal component analysis (PCA) and correlations of hormonal and inflammatory signals.** Pairwise PCA and cross-correlations of ACTH, cortisol and inflammatory mediators to determine the similarity between their dynamic profiles.

#### Supplementary Tables

**Table S1. Range of parameter values and statistical moments for optimisations of the controls and CABG groups. The hill coefficient, an integer, was fixed at 2 as the majority of fits to the controls had this value.**

| Parameter | Min | Max | Controls | | CABG | |
|---|---|---|---|---|---|---|
| | | | Median | STD | Median | STD |
| $\lambda_f\ (min)$ | 0.1 | 10 | 1.96 | 2.36 | 1.25 | 2.55 |
| $\lambda_s\ (min)$ | 1 | 1000 | 579.13 | 120.99 | 698.90 | 202.40 |
| $p_f\ (nM \cdot min^{-1})$ | 0 | 2750 | 497.00 | 681.22 | 360.77 | 682.33 |
| $p_s\ (nM \cdot min^{-1})$ | 0 | 28 | 1.49 | 0.61 | 1.24 | 0.40 |
| $K_A\ (ng \cdot L^{-1})$ | 1 | 60 | 50.28 | 9.25 | 30.19 | 5.24 |
| $m$ | 1 | 4 | 2 | N/A | 2 | N/A |

**Table S2. Range of parameter values and statistical moments for optimisations of the CABG subgroups.**

| Parameter | Two Pulses | | Multiple Pulses | | Single Pulse | |
|---|---|---|---|---|---|---|
| | Median | STD | Median | STD | Median | STD |
| $\lambda_f\ (min)$ | 1.91 | 2.78 | 0.70 | 1.56 | 1.02 | 2.58 |
| $\lambda_s\ (min)$ | 730.66 | 181.71 | 646.06 | 177.98 | 83.54 | 33.63 |
| $p_f\ (nM \cdot min^{-1})$ | 243.79 | 637.25 | 411.68 | 638.41 | 211.65 | 620.25 |
| $p_s\ (nM \cdot min^{-1})$ | 0.97 | 0.33 | 2.65 | 0.51 | 8.79 | 4.39 |
| $K_A\ (ng \cdot L^{-1})$ | 51.57 | 5.98 | 35.53 | 3.28 | 34.63 | 5.050 |
| $m$ | 2 | N/A | 2 | N/A | 2 | N/A |

**Table S3. Range of parameter values and statistical moments for optimisations of the controls and CABG groups when adrenal sensitivity is fixed.**

| Parameter | Controls | | CABG | |
|---|---|---|---|---|
| | Median | STD | Median | STD |
| $\lambda_f\ (min)$ | 2.01 | 2.38 | 1.35 | 2.60 |
| $\lambda_s\ (min)$ | 595.62 | 145.05 | 777.38 | 156.11 |
| $p_f\ (nM \cdot min^{-1})$ | 525.32 | 691.77 | 330.33 | 692.46 |
| $p_s\ (nM \cdot min^{-1})$ | 1.50 | 0.47 | 1.45 | 0.45 |
| $K_A\ (ng \cdot L^{-1})$ | 50.28 | N/A | 50.28 | N/A |
| $m$ | 2 | N/A | 2 | N/A |

**Table S4. Range of parameter values and statistical moments for optimisations of the CABG subgroups with adrenal sensitivity fixed.**

| Parameter | Two Pulses | | Multiple Pulses | | Single Pulse | |
|---|---|---|---|---|---|---|
| | Median | STD | Median | STD | Median | STD |
| $\lambda_f\ (min)$ | 1.80 | 2.72 | 0.84 | 2.17 | 1.16 | 2.57 |
| $\lambda_s\ (min)$ | 731.21 | 178.08 | 782.45 | 141.95 | 117.49 | 54.81 |
| $p_f\ (nM \cdot min^{-1})$ | 252.83 | 661.87 | 343.88 | 648.26 | 238.57 | 633.23 |
| $p_s\ (nM \cdot min^{-1})$ | 0.97 | 0.30 | 3.31 | 0.46 | 6.59 | 4.11 |
| $K_A\ (ng \cdot L^{-1})$ | 50.28 | N/A | 50.28 | N/A | 50.28 | N/A |
| $m$ | 2 | N/A | 2 | N/A | 2 | N/A |
